# Critical Nodes of Virus–Host Interaction Revealed Through an Integrated Network Analysis

**DOI:** 10.1101/548909

**Authors:** Korbinian Bösl, Aleksandr Ianevski, Thoa T. Than, Petter I. Andersen, Suvi Kuivanen, Mona Teppor, Eva Zusinaite, Uga Dumpis, Astra Vitkauskiene, Rebecca J. Cox, Hannimari Kallio-Kokko, Anders Bergqvist, Tanel Tenson, Valentyn Oksenych, Magnar Bjørås, Marit W. Anthonsen, David Shum, Mari Kaarbø, Olli Vapalahti, Marc P. Windisch, Giulio Superti-Furga, Berend Snijder, Denis Kainov, Richard K. Kandasamy

## Abstract

Viruses are one of the major causes of various acute and chronic infectious diseases and thus a major contributor to the global burden of disease. Several studies have shown how viruses have evolved to hijack basic cellular pathways and evade innate immune response by modulating key host factors and signalling pathways. A collective view of these multiple studies could advance our understanding of viral evasion mechanisms and provide new therapeutic perspectives for the treatment of viral diseases. Here, we performed an integrative meta-analysis to elucidate the 17 different host-virus interactomes. Network and bioinformatics analyses showed how viruses with small genomes efficiently achieve the maximal effect by targeting multifunctional and highly connected host proteins with a high occurrence of disordered regions. We also identified the core cellular process subnetworks that are targeted by all the viruses. Integration with functional RNA interference (RNAi) datasets showed that a large proportion of the targets are required for viral replication. Furthermore, we performed an interactome-informed drug re-purposing screen and identified novel activities for broad-spectrum antiviral agents against hepatitis C virus and human metapneumovirus. Altogether, these orthogonal datasets could serve as a platform for hypothesis generation and follow-up studies to broaden our understanding of the viral evasion landscape.

## Introduction

Viruses continue to be a major contributor to the global burden of disease through acute and chronic infections that cause substantial economic impact in addition to increased mortality and morbidity (Virgin et al. 2009). Despite the tremendous improvement in the understanding of the antiviral immune response and the availability of therapeutics, existing and emerging viral diseases are an ever-growing problem, particularly in developing countries. Development of antiviral resistance of viruses such as hepatitis C virus (HCV), influenza A virus (IAV), herpes simplex virus (HSV), human cytomegalovirus (HCMV) and human immunodeficiency virus (HIV) is a major concern (Irwin et al. 2016, Howard and Fletcher 2012, Bacon et al. 2003). One of the main reasons for increasing resistances is the accumulation of mutations in the viral genome caused by multiple factors including the polymerase infidelity (Sanjuan and Domingo-Calap 2016, Peck and Lauring 2018). Therefore, the World Health Organization (WHO) has urged for better control of viral diseases.

This has led to turning the focus on the host for therapeutic intervention, where targeting the host factors has been proven to be useful for restricting viral infections (Zumla et al. 2016, Kaufmann et al. 2018). Anti-CXCR4 and anti-CCR5 monoclonal antibodies are examples of successful host-directed therapies for combating HIV (Olson and Jacobson 2009, Carnec et al. 2005).

Viruses have evolved to evade the host antiviral response at various stages starting from viral sensing to antiviral pro-inflammatory responses (Bowie and Unterholzner 2008, Navratil et al. 2011, Garcia-Sastre 2017). Multiple studies attempted to understand global principles of the viral evasion employed by various viruses, including dengue virus (DENV), Ebola virus (EBOV), IAV and HIV (Pichlmair et al. 2012, Rozenblatt-Rosen et al. 2012, Jäger et al. 2011, Shah et al. 2018, Batra et al. 2018, Söderholm et al. 2016). Global systems-level approaches including functional RNAi screens, interactome mapping technologies such as affinity-purification mass spectrometry (AP-MS), quantitative proteomics and CRISPR/Cas9-based screens have provided unparalleled details and insights into the dynamics of host proteome in immune cells (Weekes et al. 2014, Nightingale et al. 2018, Kandasamy et al. 2016, Ersing et al. 2017), host-virus interactome (Pichlmair et al. 2012, Rozenblatt-Rosen et al. 2012, Jäger et al. 2011, Shapira et al. 2009, Konig et al. 2010), and also identified important host dependency factors of various viruses (Shapira et al. 2009, Zhang et al. 2016, Marceau et al. 2016). Meta-analyses of such high-dimensional datasets have been crucial for identifying novel host factors as drug targets such as UBR4 in IAV infection (Tripathi et al. 2015). Moreover, some of these factors represent drug targets for multiple viruses(Ianevski et al. 2018).

We hypothesized that combining a meta-analysis of host-virus protein-protein interactions of multiple viruses and functional RNAi screens would provide novel insights for developing broad-spectrum antiviral strategies. For this, we assembled a host-virus protein-protein interactome of 5781 host-virus interactions (hereafter referred to as ‘hvPPI’) covering 183 viral proteins from 17 different viruses and 2381 host proteins. We performed extensive bioinformatics and network analysis and integrated this dataset with genome-wide or druggable-genome RNAi screen data from published studies. This resulted in the assembly of critical nodes of viral evasion and identification of core cellular processes and druggable nodes that were verified by a drug re-purposing screen using broad-spectrum antivirals.

## Material & Methods

### Construction of hvPPI data, network analysis and data visualization

Host-virus protein-protein interactions were downloaded from published studies (Pichlmair et al. 2012, Rozenblatt-Rosen et al. 2012, Jäger et al. 2011, Shapira et al. 2009, de Chassey et al. 2008, Khadka et al. 2011, Zhang et al. 2009, Vidal et al. 2011) which included a total of 183 viral proteins, 2381 host proteins and 5781 host-virus interactions. Protein identifiers were mapped to UniProt IDs. Human protein-protein interaction data was imported from BioGRID database (version 3.4.139, Stark et al. 2006) covering 215244 interactions. The network analysis was performed using in-house programs developed in R statistical environment (version 3.4.3, www.r-project.org) with the use of the packages SparseM (version 1.77), RBGL (version 1.52.0) and graph (version 1.54.0). Network visualization was performed in Cytoscape (version 3.6.1, Shannon et al. 2003). Network clusters/sub-networks were extracted using the Cytoscape plugin MCODE (version 1.5.1, Bader and Hogue 2003). Data visualization was performed in R statistical environment and Cytoscape.

### Gene-set enrichment, protein disorder and sub-cellular localization analysis

We performed gene-set enrichment analysis using DAVID Bioinformatics Resources (version 6.8, Huang da et al. 2009). For all enrichment analysis, a p-value cutoff of ≤0.01 was used as significant. Protein disorder analysis was performed using IUPred2A software. We used the offline version with protein sequences downloaded from UniProt. Statistical analysis of disordered region distribution was performed by Kolmogorov-Smirnov test in R statistical environment. Annotation of human proteins was mapped from UniProt ID to ENSEMBL using EnsDb.Hsapiens.v86. The index of subcellular localization of interaction partners of single viral proteins was calculated for all viral proteins with ≥ five host targets. Localization of host targets was mapped using COMPARTMENTS (Binder et al. 2014), filtered for a minimum evidence score of 3 in the knowledge channel, excluding non-experimental based localization predictions. Evidence for all protein was subsequently divided by the absolute number of host-targets per viral protein. Multiple sequence alignment was performed using Clustal X (version 2.0, Larkin et al. 2007).

### Integration of RNAi screens and drug-gene interaction data

Genome-wide RNAi screen data for HCV (Tai et al. 2009) and HPV18 (Smith et al. 2010, through GenomeRNAi database (Schmidt et al. 2013- GR00197), as well as druggable RNAi screen data for HPV16 (Aydin et al. 2014), VACV (Mercer et al. 2012) and SV40 (Snijder et al. 2012) were integrated in the existing network as Z-Scores. Drug-gene interaction data was downloaded from DGIdb. The identifiers were mapped to UniProt IDs and then compared with hvPPI.

### Drug re-purposing screen

For the HMPV NL/1/00 screen, approximately 4×104 human retinal pigment epithelial (RPE) cells were seeded per well in 96-well plates. The cells were grown for 24 h in DMEM-F12 medium supplemented with 10% FBS, 0.35% NaHCO3, and 100 μg/ml streptomicine and 100 IU/ml penicillin. The medium was replaced with virus growth medium (VGM) containing 0.2% bovine serum albumin (BSA), 2 mM L-glutamine, 0.35% NaHCO3, and 1 μg/mL L-1-tosylamido-2-phenylethyl chloromethyl ketone-trypsin in DMEM-F12. HCV screen-associated cell culture conditions are described in Kim et al. (2016). The compounds were added to the cells in 3-fold dilutions at seven different concentrations starting from 50 μM. No compounds were added to the control wells. The cells were mock- or virus-infected at a multiplicity of infection (MOI) of one. After 48 h of infection, the medium was removed from the cells. To monitor cell viability, CellTiter-Glo reagent containing Firefly luciferase and luciferin was added (30 μL per well). This assay quantifies ATP, an indicator of metabolically active living cells. The luminescence was measured with a plate reader. To determine compound efficacy, HMPV NL/1/00-induced GFP expression was measured. The half-maximal cytotoxic concentration (*CC*_50_) and the half-maximal effective concentration (*EC*_50_) for each compound were calculated after non-linear regression analysis with a variable slope using GraphPad Prism software version 7.0a. The relative effectiveness of the drug was quantified as the selectivity index 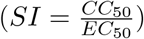.

Cytotoxicity and antiviral activity of the compounds against GFP-expressing HCV in Huh-7.5 cells was determined as previously described (Kim et al. 2016).

## Results

### Assembly of host-virus protein-protein interactions

To provide new and critical insights into viral evasion mechanisms we performed a comprehensive meta-analysis of the host-virus interaction landscape. We assembled the host-virus protein-protein interaction data (‘hvPPI’) from published studies (Fig. 1A) (Pichlmair et al. 2012, Rozenblatt-Rosen et al. 2012, Jäger et al. 2011, Shapira et al. 2009, de Chassey et al. 2008, Khadka et al. 2011, Zhang et al. 2009, Vidal et al. 2011). This dataset covered 17 different viruses including adeno-associated virus 5 (AAV5), dengue virus (DENV), Epstein-Barr virus (EBV), influenza A virus PR8 (IAV-PR8), influenza virus Udorn (IAV-Udorn), hepatitis C virus (HCV), human immunodeficiency virus 1 (HIV-1), human papilloma virus 5 (HPV5), human papilloma virus 6B (HPV6B), human papilloma virus 8 (HPV8), human papilloma virus 11 (HPV11), human papilloma virus 16 (HPV16), human papilloma virus 18 (HPV18), human papilloma virus 33 (HPV33), Merkel cell polyomavirus (MCPyV), Simian virus 40 (SV40) and Vaccinia virus (VACV). This dataset comprised of protein-protein interactions from two different types of experimental methods - affinity purification mass spectrometry (AP-MS) and yeast two-hybrid screens (Y2H). Altogether, this combined dataset includes 183 viral proteins, 2381 host proteins and 5781 protein-protein interactions (Fig. 1B and Fig. S1). Many interactome networks including yeast and human are scale-free networks, where a large portion of the nodes (e.g. a protein in the network) have few interactions and only a few nodes have large number of interactions. The latter are often referred to as “hubs” which are crucial in keeping the network intact (Vidal et al. 2011). We performed network topology analysis to infer the properties of the host proteins targeted by the viral proteins in the context of the human protein interactome. We considered two important parameters - relative betweenness centrality (which reflects the amount of information that passes through this protein in the human interactome) and degree (number of binding partners in the human interactome) of the host proteins targeted by each virus. The targets of all the viruses showed higher betweenness centrality and degree as compared to an average protein in the human interactome (Fig. 1C and Fig. 1D). This shows that viruses, by targeting “hubs” and proteins that serves as key communication nodes, have evolved the best way to disrupt the scale-free human interactome. This topological property thereby shows how viruses having small genomes achieve the maximal effect in rewiring the human interactome to benefit viral survival and replication. Our analysis is in agreement with several previous studies, which have highlighted this property (Pichlmair et al. 2012, Rozenblatt-Rosen et al. 2012, de Chassey et al. 2008, Durmuş et al. 2015, Franzosa and Xia 2011). We propose that this could be a general principle for all viruses.

**Fig. 1:**
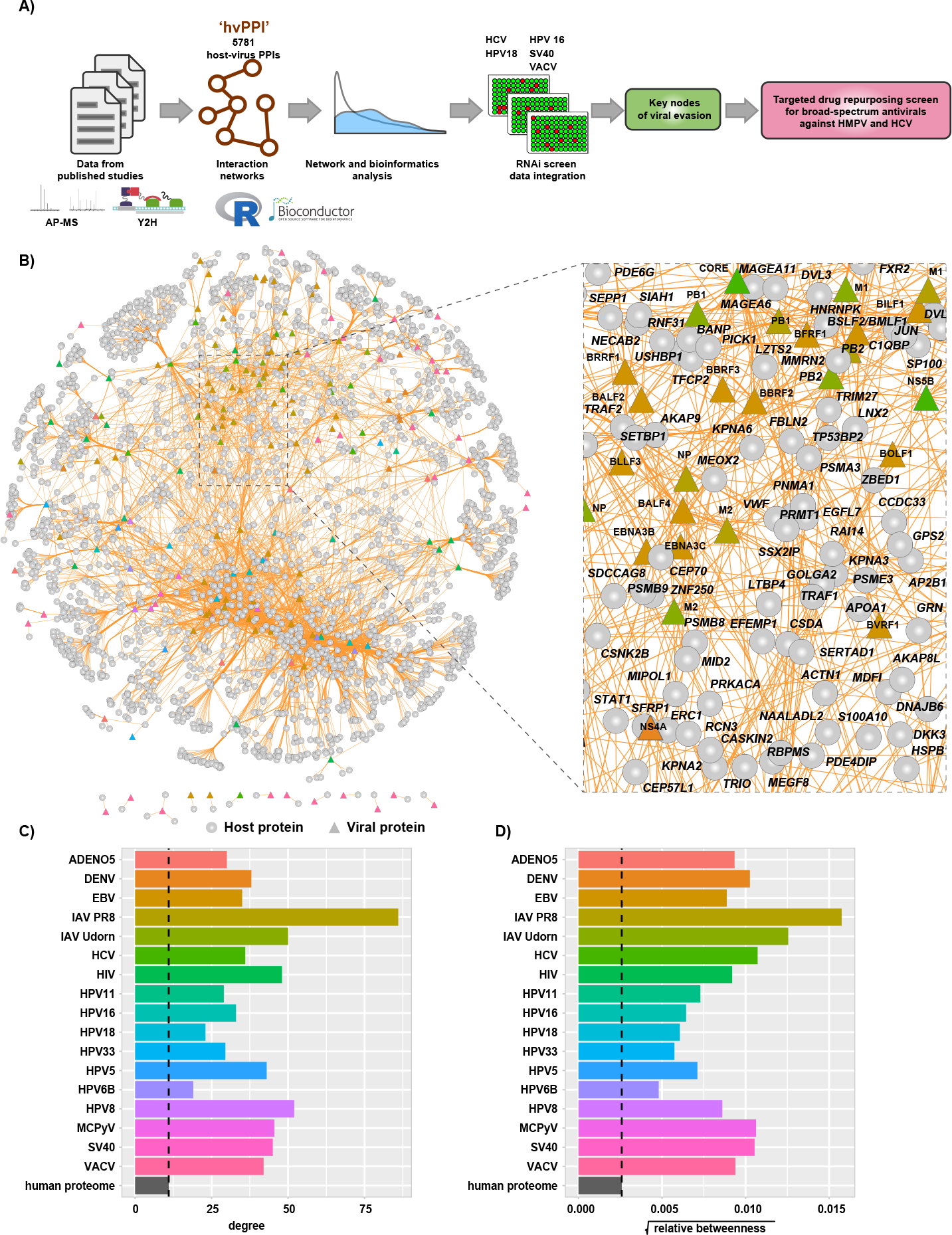
Meta-analysis of host-virus interactions involving 17 different viruses. **(A)** Schematic view of the analysis workflow. **(B)** Network view of the ‘hvPPI’ containing host-virus interactions from 17 different viruses. The edges are coloured in orange. Node shapes are in circles and triangles for host and viral protein, respectively. A zoomed-in snippet shows the names of selected host and viral proteins. **(C)** and **(D)** Barplot showing the median degree and betweenness centrality of targets of each virus as compared to the human proteome.

### Host factors with higher disordered regions are enriched in hvPPI networks

Proteins usually fold into stable three-dimensional structures that mediate specific functions. In addition, there are sub-structures in proteins termed “intrinsically disordered regions (IDRs)” which lack stable structures under normal physiological conditions. IDRs are required for multiple cellular functions even though they lack these defined structures (Oldfield and Dunker 2014). Many studies have highlighted the presence of such IDRs in viral proteins (Xue et al. 2014, Tamarozzi and Giuliatti 2018, Dyson and Wright 2018), such as E6 from human papilloma virus, that are crucial for hijacking the cellular machinery. We analysed the host proteins from the hvPPI for presence of IDRs using the prediction software IUPred (Meszaros et al. 2018). We found a statistically significant enrichment (p-value < 6.246 × 10^−06^) of IDRs in the host proteins targeted by viruses (Fig. 2A and Fig. S2A). We then identified the subnetwork in the hvPPI which contained the top host targets with high disorderness score (Fig. 2B). The top five proteins with large IDRs include CD44 antigen (CD44), Serine/arginine repetitive matrix protein 2 (SRRM2), Myristoylated alanine-rich C-kinase substrate (MARCKS), BAG family molecular chaperone regulator 3 (BAG3) and Mitochondrial antiviral-signalling protein (MAVS) (Fig. 2C). CD44 is a marker of exhausted CD8+ T cells (Wherry and Kurachi 2015) and replication of HCV in T cells was shown to decrease cell proliferation by inhibiting CD44 expression and signalling (Kondo et al. 2009). SRRM2 is a serine/arginine-rich protein involved in RNA splicing (Blencowe et al. 2000). SRRM2 is differentially phosphorylated in HIV-1 infected cells and absence of SRRM2 lead to increased HIV-1 gene expression, since it regulates the splicing of HIV-1 (Wojcechowskyj et al. 2013). In the hvPPI, SRRM2 is targeted by multiple viral proteins including the Tat protein from HIV-1. Tat protein has an important role in the stimulation of the transcription of the long terminal repeat (LTR) (Das et al. 2011). In addition, NS1 protein from influenza B virus has also been reported to interact with SRRM2 (Patzina et al. 2017). Proteins of the MARCKS family are involved in a range of cellular processes including cell adhesion and migration (Arbuzova et al. 2002). MARCKS is a negative regulator of lipopolysaccharide (LPS)-induced Toll-like receptor 4 (TLR4) signalling in mouse macrophages (Mancek-Keber et al. 2012). MAVS is an adaptor protein in the RIG-I signalling pathway involved in the sensing of RNA. Ablasser et al. (2009) reported that doublestranded DNA serves as a template for RNA polymerase II and is transcribed into a 5’ triphosphate containing double-stranded RNA, which activates the RIG-I signalling pathway. In the hvPPI, MAVS is targeted by several proteins from dsDNA viruses such as EBV and HPV. Altogether, our analysis shows that the IDR-high part of the human proteome is an essential part of the viral evasion strategy and some of the selected targets highlighted here could show novel insights into the viral evasion mechanisms. However, the very flexible protein structure of disordered proteins also makes them also difficult to target with drugs.

**Fig. 2:**
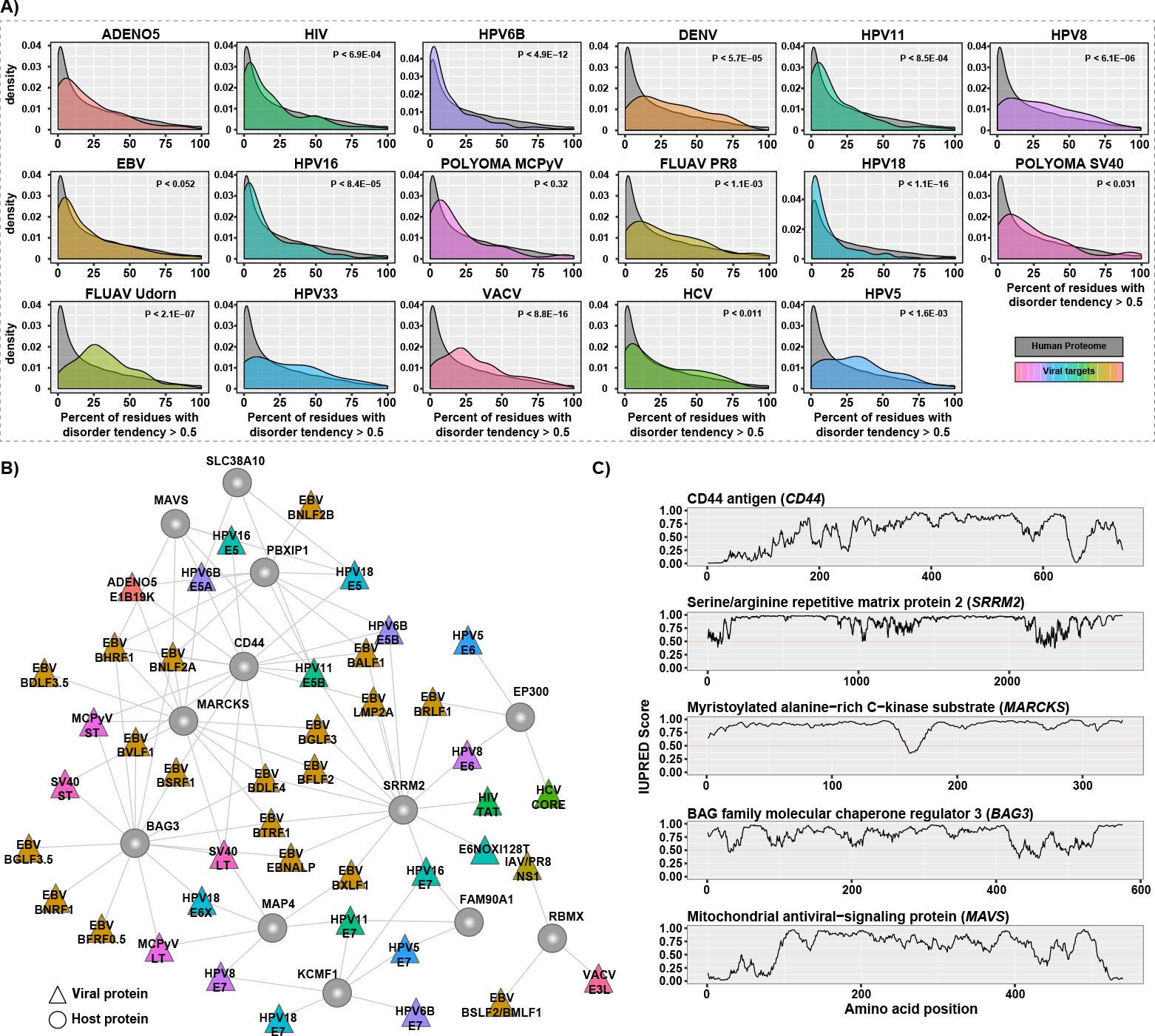
Protein disorder analysis. **(A)** Density plot of the distribution of host proteins for each virus with percent of residues with disorder tendency greater than 0.5 as predicted by the IUPred software as compared to the human proteome. **(B)** Sub-network of hvPPI with the highly targeted and highly-disordered proteins. **(C)** Line plots showing the IUPred Score (a measure of the disordered region) for the five selected host proteins from the sub-network. A IUPred score of >0.5 is considered disordered.

### Viral proteins target core signalling pathways and process networks

To assess the signalling pathways and cellular processes within the hvPPI, we identified highly connected subnetworks within hvPPI network. We constructed a host-host interaction network based on the host targets in the hvPPI and identified a number of highly connected subnetworks/clusters (Fig. 3). We then performed a gene-set enrichment analysis of significantly enriched biological processes. We found one or more enriched processes for each of this subnetwork including core cellular processes such as proteasome, spliceosome, protein translation, protein/RNA transport, and cell cycle. Next we listed the viruses that target one or more of these processes, and found that almost all the core pathways and processes are targeted by all the 17 viruses that are part of the hvPPI (Fig. S3). This analysis highlights the core components of the cellular process subnetworks which are targeted as part of the viral evasion strategies and thus could be broad-spectrum antiviral hot-spots from a therapeutic point of view.

**Fig. 3:**
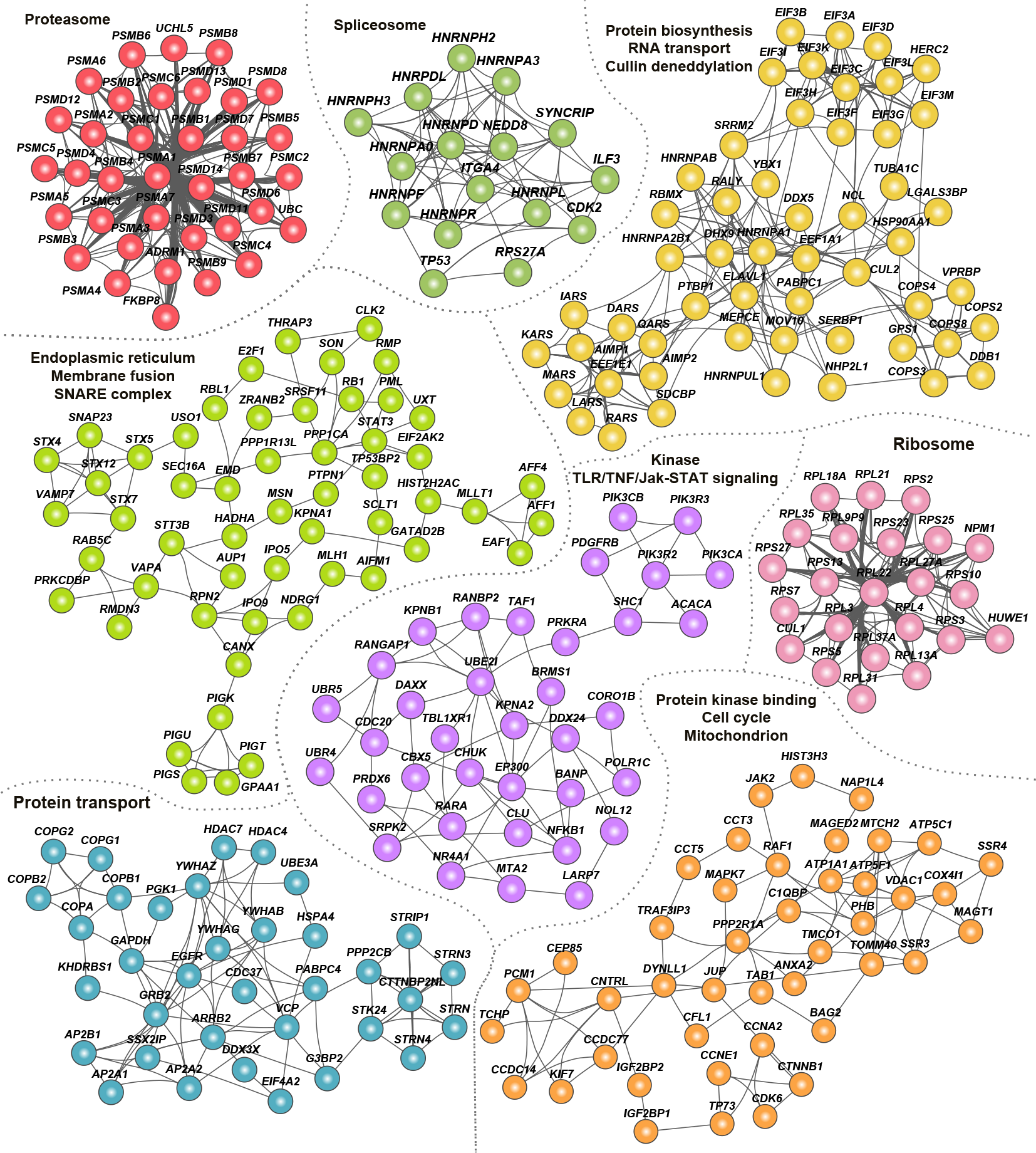
Clusters of hvPPI involved in core cellular processes. Network view of the ‘clusters’ or highly-connected sub-networks and their associated cellular processes. Each cluster is marked in a unique color.

### Enrichment analysis reveals commonality and specificity in sub-cellular localization of the host factors

Given that the viral proteins were interacting with a large number of host proteins, we analysed the sub-cellular location of the host proteins. We performed gene-set enrichment analysis of sub-cellular localization information provided by UniProt database. We binned the localization into 11 compartments and estimated the percent of host proteins in a given compartment as compared to the total number of host proteins targeted by a given virus. We found that the viral targets were distributed across multiple subcellular compartments with cytoplasm being the most common (Fig. S4A). The hvPPI includes two different strains of IAV-PR8 (H1N1) and Udorn (H3N2). The subcellular localization analysis showed that both strains were enriched for nuclear proteins. Nonstructural protein 1 (NS1) from both the strains had the highest number of nuclear targets but their targets were very different (Fig. 4A). NS1 of Udorn was enriched for a large number of histones as compared to NS1 of PR8 that had large number of heterogeneous nuclear ribonucleoproteins (hnRNPs), such as HNRNPU - a known restriction factor for many viruses. This corroborates with the observation that NS1 protein has short linear histone mimicry motifs that can suppress the host antiviral response (Marazzi et al. 2012). In our analysis, we found that it is NS1 of Udorn that has a histone mimicry motif “ARSK” (Fig. S4B). Similarly, HPV11 and HPV18 E5 proteins interact more often with host proteins located in the endoplasmic reticulum (ER). We found both common and specific subsets of ER proteins targeted by the E5 protein (Fig. 4B). HPV18 E5 protein ER targets were enriched for phospholipid biosynthesis as well as GPI anchor related proteins, such as phosphatidylinositol glycan anchor biosynthesis class S/T/U (PIGS, PIGT and PIGU), glycosylphosphatidylinositol anchor attachment 1 (GPAA1) and phosphatidylserine synthase 2 (PTDSS2). HPV11 E5 protein ER targets were enriched for ER-associated ubiquitin-dependent protein catabolism involving host proteins such as ER degradation enhancing alpha-mannosidase-like protein 3 (EDEM3) and ER lipid raft associated 1 (ERLIN1). ER targets common to HPV18 and HPV11 E5 protein were enriched for unfolded protein response, N-linked glycosylation and protein folding involving host proteins such as SRP receptor alpha/beta subunit (SRPRA/SRPRB) and catalytic subunits of the oligosaccharyltransferase complex (STT3A and STT3B). Two independent CRISPR/Cas9 screening studies identified multiple ER associated components including STT3A and STT3B as host factors for DENV, Zika virus (ZIKV) and Japanese encephalitis virus (JEV) (Zhang et al. 2016, Marceau et al. 2016). The non-canonical function of STT3A and STT3B is required for DENV replication and that NS1 protein of DENV interacts with these proteins (Marceau et al. 2016). Our orthogonal approach can lead to the identification of critical host factors, and similar functions of ER components, such as STT3A and STT3B, are used by HPV11 and HPV18 as well. Thus targeting the non-canonical function of STT3A and STT3B could be a broad antiviral strategy. Overall, the enrichment analysis clearly shows that there is commonality and specificity in the subcellular targets of the viral proteins and that detailed interrogation of these targets can give vital clues into the viral evasion mechanisms.

**Fig. 4:**
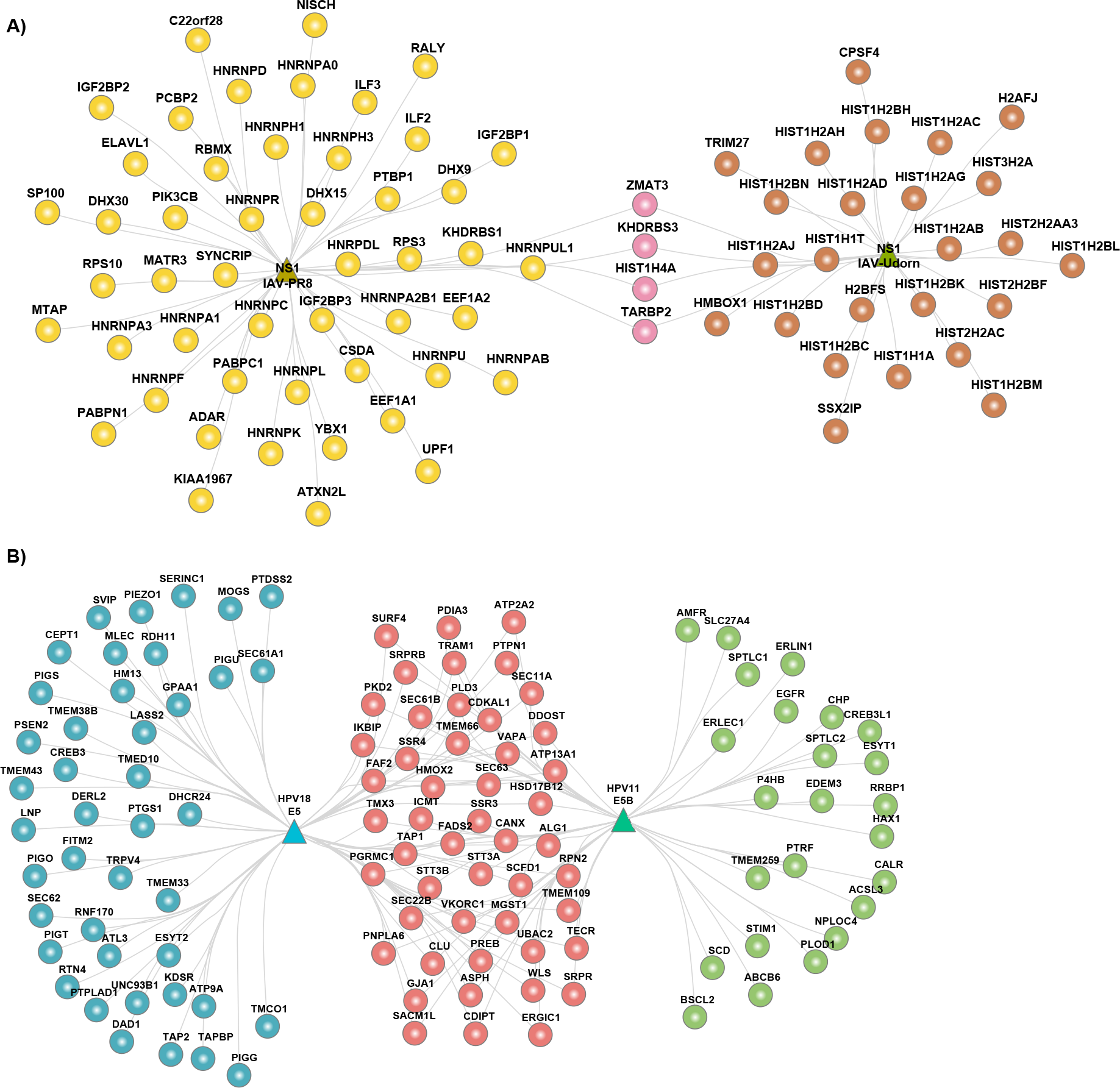
Sub-cellular localization of the host proteins. **(A)** Network view of nuclear interactome of NS1 protein from IAV strains PR8 and Udorn. **(B)** Network view of ER interactome of E5 protein from HPV18 and HPV11.

### Integrative analysis of host-virus interactome and RNAi data reveals COPI system as commonly targeted proviral process

RNAi screens have been a powerful high-throughput method to identify various cellular functions, including for identification of host restriction factors of viruses (Hirsch 2010). In order to explore the functional relevance of the host targets in the hvPPI, we integrated it with five published RNAi screens that performed genome-wide or druggable-genome-wide RNAi screens for identifying host factors of HCV (Tai et al. 2009), HPV18 (Schmidt et al. 2013), HPV16 (Aydin et al. 2014), SV40 (Snijder et al. 2012) and VACV (Mercer et al. 2012).

We found that host targets from the hvPPI were spread across the spectrum of genes with proviral as well as antiviral phenotype (Fig. S5), thus showing that targeting of the host protein by the virus could lead into any direction that favours the virus. We then investigated the top 50 proviral genes that are also targeted by the viral proteins as seen in the hvPPI. We identified 42 host proteins (Fig. 5A) that were significantly enriched for coatomer protein complex 1 (COPI), protein translation/transport and proteasome (Fig. 5B). This further substantiates the findings from the earlier section on the core cellular processes targeted by the viruses. Network analysis of these top hits showed high level on connectivity and crosstalk - for example between the translation and proteasome machinery (Fig. 5C). Vesicle carriers are involved in the transport of membranes and proteins. COPI system is one of the three vesicular carrier systems that is involved in the early secretory pathway (Beck et al. 2009, Lee et al. 2004, Bethune et al. 2006). Moreover, it has been pointed out that there is a strong similarity between vesicular transport and viral transport (viral entry to budding process) (Thaa et al. 2010); thus making COPI system important for the viral life cycle. SiRNA-based silencing of COPI lead to a defect in entry of IAV and disruption of the COPI complex inhibited the production of infectious progeny virus (Sun et al. 2013). Altogether, these top hits including the COPI system could serve as targets for developing therapeutic antiviral intervention strategies for a broad group of viruses.

**Fig. 5:**
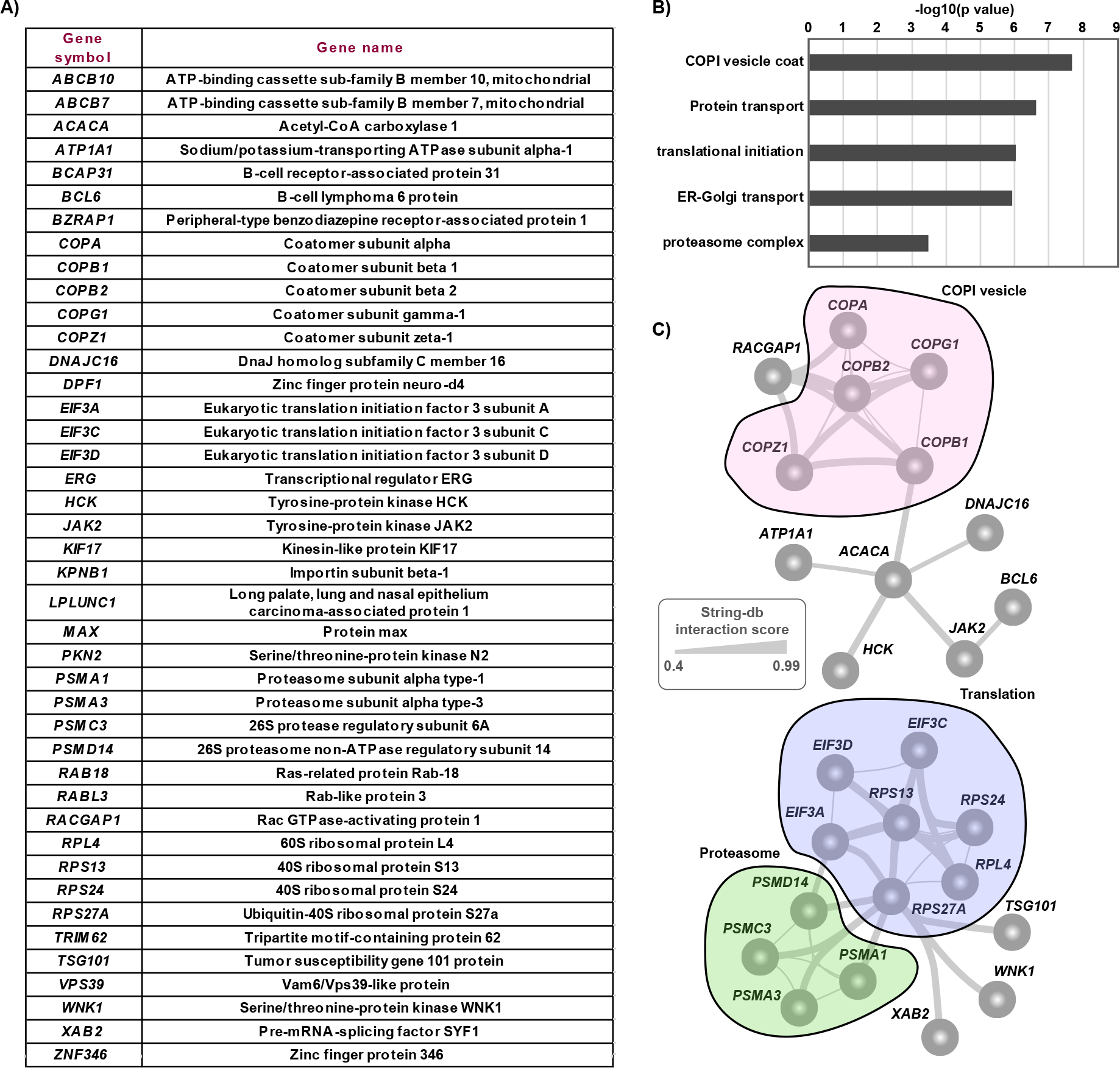
Integration of hvPPI with RNAi screen. **(A)** Top proviral genes from RNAi screens that are also targeted by multiple viral proteins. **(B)** Barplot showing the significantly enriched cellular processes involving the top targeted and proviral genes. **(C)** Network view of top targets and their functional relevance.

### Combining host-virus and drug-gene interactions reveals novel activities of broad-spectrum antiviral agents against hepatitis C virus and human metapneumovirus

Our analyses pointed out that viral evasion mechanism observed in one virus could also be relevant for other viruses. Thus, we wanted to test if we could target viral replication of other viruses based on the results from the hvPPI analyses. To get an estimate of the druggability of the hvPPI, we obtained known drug-gene interactions from DGIdb (Cotto et al. 2018). The data from DGIdb included drug-gene interactions for 48 drugs that were investigational/experimental/approved safe-in-human antivirals compounds (Ianevski et al. 2018). Among these 48 broad-spectrum-antivirals, 28 of them had targets that are part of the hvPPI. Based on this overlap, we performed a targeted drug re-purposing screen with 35 of these 48 compounds (Table S1) to check for antiviral activity against and HCV, which is part of the hvPPI and human metapneumovirus (HMPV) which is not in the hvPPI.. We tested 35 broad-spectrum-antivirals against GFP-expressing HMPV NL/1/00 strain (de Graaf et al. 2007). Seven different concentrations of the compounds were added to HMPV or mock-infected cells. Cell viability was controlled after 48 hours and HMPV-induced GFP expression was tested to determine compound efficiency. After the initial screening, we identified five compounds, which lowered GFP-expression without detectable cytotoxicity (with *SI* > 3). We repeated the experiment with these compounds. The experiment confirmed novel activity of azacytidine, lopinavir, nitazoxanide, itraconazole and oritavancin against HMPV (Fig. 6A and Table 1). Similarly, we examined toxicity and antiviral activity of broad-spectrum-antivirals against GFP-expressing HCV in Huh-7.5 cells using previously described procedures (Kim et al. 2016). Our test identified azithromycin, cidofovir, oritavancin, dibucaine, gefitinib, minocycline and pirlindole mesylate as novel anti-HCV agents with *SI* > 3 (Fig. 6B and Table 1).

**Fig. 6:**
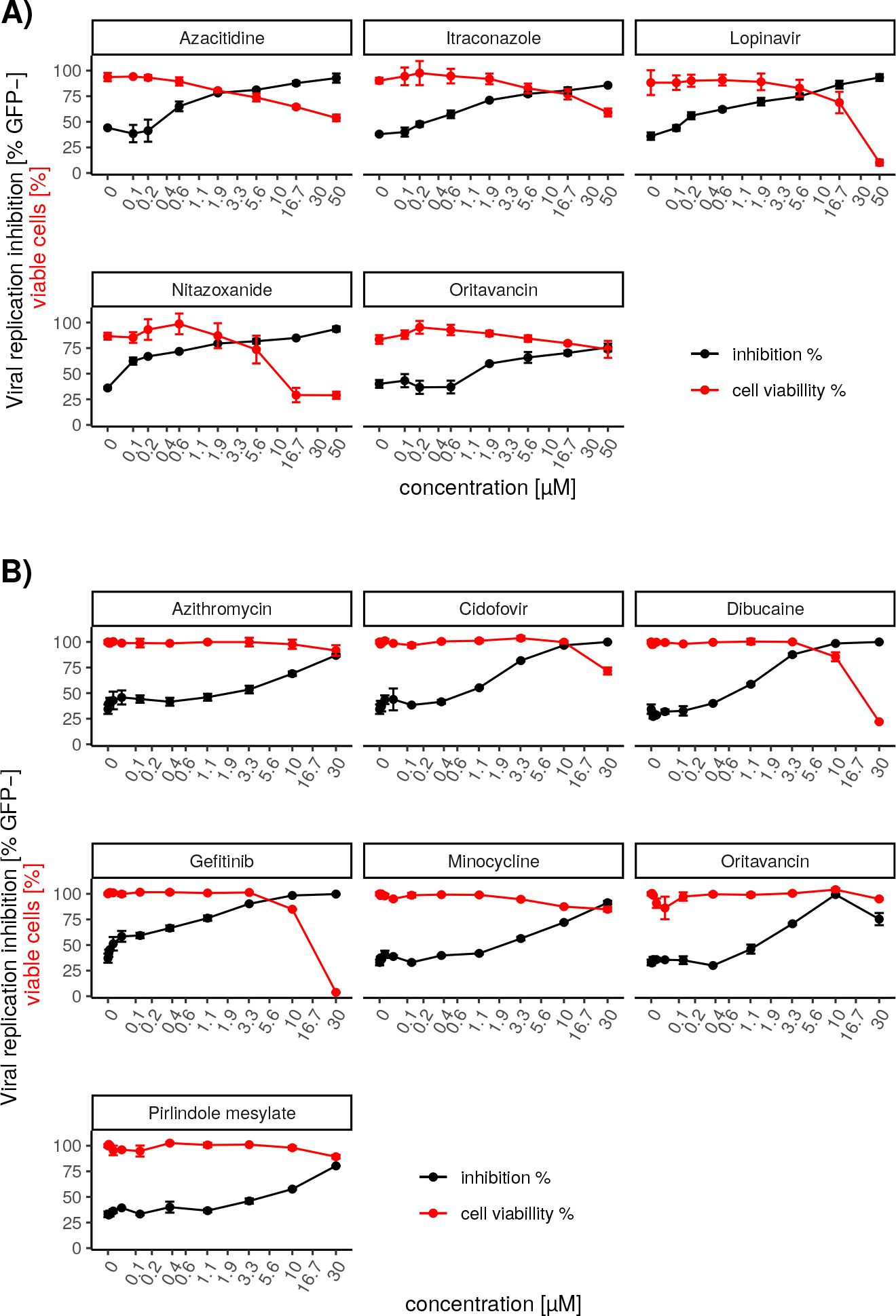
Drug re-purposing screen for novel broad-spectrum antivirals **(A)** At non-cytotoxic concentrations selected broad-spectrum antiviral agents inhibit HMPV-mediated GFP expression in Huh-7.5 cells. Efficacy of antiviral agents as shown by the percent of GFP-expressing cells decreasing in response to increasing concentration of antiviral agents (Mean ±SD, n=3). Cytotoxicity of antiviral agents as shown by the percent of live cells (Mean ±SD, n=3). **(B)** At non-cytotoxic concentrations selected broad-spectrum antiviral agents inhibit HCV-mediated GFP expression in Huh7.5 cells (Mean ±SD, n=3).

**Table 1:**
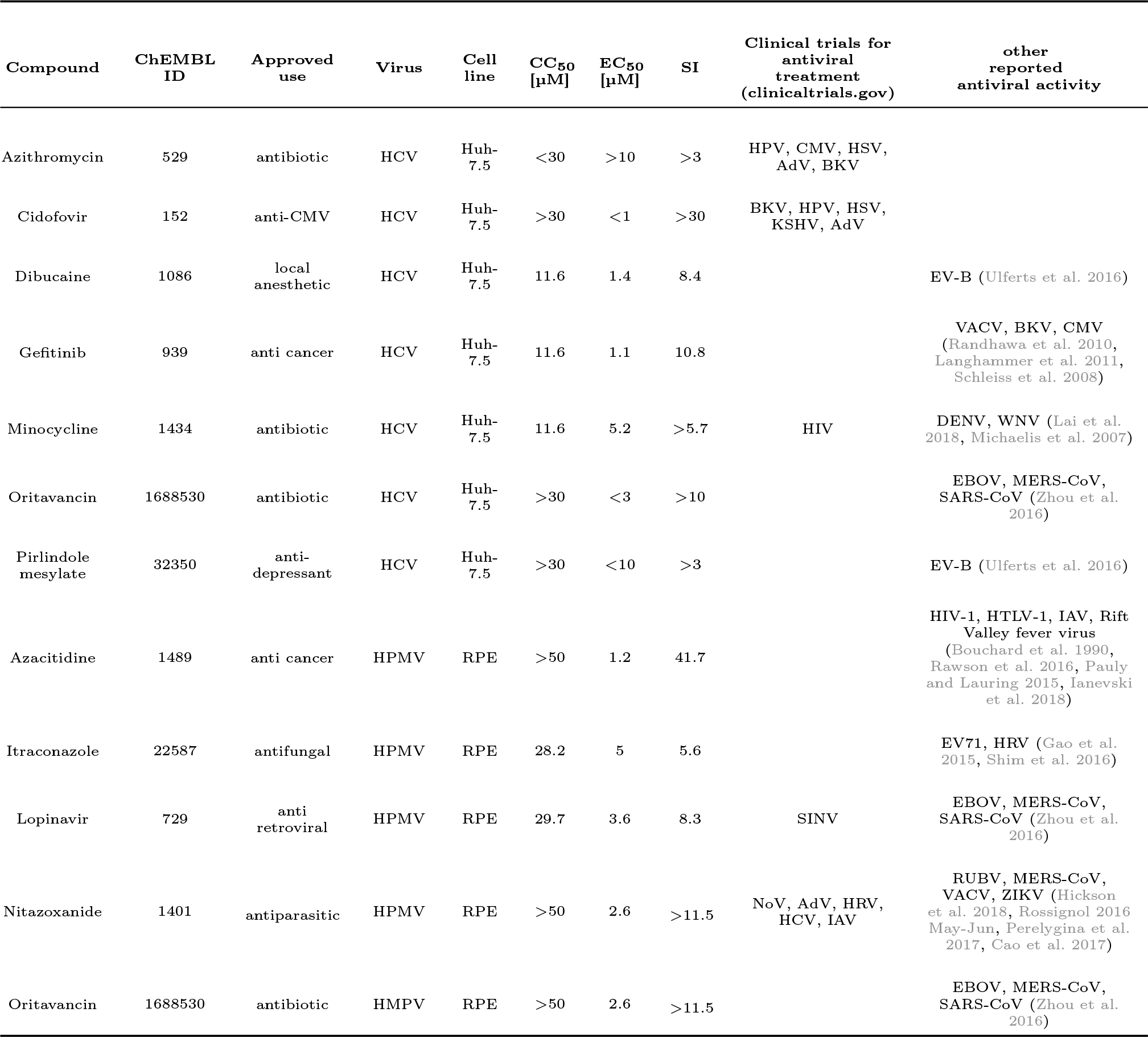
Characteristics, half-maximal cytotoxic concentration (*CC*_50_), the half-maximal effective concentration (*EC*_50_) and minimal selectivity indexes 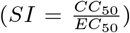 for selected broad-spectrum antivirals. The measurements were repeated three times (p<0.05).

In summary, our meta-analysis approach of the hvPPI could provide novel and faster approaches for the re-purposing of existing drugs as antiviral agents.

## Discussion

Using integrative analysis of orthogonal datasets our study provides a comprehensive view of viral evasion mechanisms.

In particular our analysis of the hvPPI network revealed that all the viruses have evolved to target proteins that are central and have strong control over the human interactome. Host proteins targeted by viruses contain a high proportion of intrinsically disordered regions. We identified the core cellular processes and associated proteins that are targeted by all viruses. Detailed comparative analysis of the subcellular localization of the host proteins showed commonality and specificity both between viral proteins from different strains of the same virus; and between viruses. Integrating hvPPI with functional RNAi screens showed that a large portion of the hvPPI are host factors of one or more virus. hvPPI data-based drug re-purposing screen identified novel activities for various broad-spectrum antivirals against HMPV and HCV.

This unique dataset can be used for further detailed interrogation of the mechanisms behind viral evasion. This could serve as a starting point for identifying novel host targets and generating hypothesis in the context of viral evasion and development of pan-viral therapeutic intervention strategies. The methods described here also provide unique ways of dissecting the orthogonal datasets. Various analyses from this study have highlighted the existence on pan-viral evasion points that could be utilised for the development of host-directed antiviral therapies. It is also intriguing to see that there is commonality and specificity at the level of subcellular localization of the viral targets. Our analyses have underlined some salient features in the context of IAV, HPV, DENV and HCV. Further detailed analysis in this context along with protein sequence features, such as Short Linear Motifs (SLiMs) (Davey et al. 2011) would provide novel insights as well as deeper understanding of how small sequence features are involved in the hijacking of the host machinery. Integration of such data with known drug-gene interactions provides a clear estimate of the druggable proportion in the hvPPI. Our meta-analysis approach of the hvPPI could provide novel avenues of re-purposing existing drugs for antiviral targeting strategies.

Our study demonstrated that azacytidine, itraconazole, lopinavir, nitazoxanide, and oritavancin are novel experimental anti-HMPV agents, whereas cidofovir, dibucaine, azithromycin, gefitinib, minocycline, oritavancin, and pirlindole mesylate are novel anti-HCV drugs. These drugs have already been used as investigational agents or experimental drug in different virus infections. Although, the mechanisms of action of these compounds in inhibiting the viral infections are still unknown, this agent could inhibit steps of viral infections, which preceded reporter protein expression from viral RNA. In summary, our results indicate that existing broad-spectrum-antivirals could be re-purposed to other viral infections. Re-purposing these therapeutics could save resources and time needed for development of novel drugs to quickly address unmet medical needs, because safety profiles of these agents are available. Effective treatment with broad-spectrum-antivirals may shortly become available, pending the results of further pre-clinical studies and clinical trials. This, in turn, means that some broad-spectrum-antivirals could be used for rapid management of new or emerging drug-resistant strains, as well as for first-line treatment or for prophylaxis of acute virus infections or for viral co-infections. The most effective and tolerable compounds could expand the available therapeutics for the treatment of viral diseases, improving preparedness and the protection of the general population from viral epidemics and pandemics.

## Conflict of Interest Statement

The authors declare that the research was conducted in the absence of any commercial or financial relationships that could be construed as a potential conflict of interest.

## Author Contributions

R.K.K. and K.B. performed all the bioinformatics and network analysis. A.I., T.T.T., P.I.A, M.T., E.Z., U.D., A.V., R.J.C., H.K-K., A.B., T.T., A.M., V.O., M.B., M.W.A., D.S., M.K., M.P.W. and D.K. contributed to the drug re-purposing screen. D.K. supervised the drug re-purposing screen. G.S.F. and B.S. provided data. R.K.K. conceived and supervised the study. R.K.K., D.K. and K.B. wrote the manuscript. All authors contributed, read and approved the manuscript.

## Funding

This work was funded by the Research Council of Norway (FRIMEDBIO “Young Research Talent” Grant 263168 to R.K.K.; and Centres of Excellence Funding Scheme Project 223255/F50 to CEMIR), Onsager fellowship from NTNU (to R.K.K.). European Regional Development Fund, the Mobilitas Pluss Project MOBTT39 (to D.K.) and by the National Research Foundation of Korea (NRF) grant funded by the Korea government (MSIT, NRF-2017M3A9G6068246, Gyeonggi-do to M.P.W.).

## Acknowledgments

We thank Christian Sinzger and group for the EGFP-expressing TB40E CMV strain. We thank ViroNovative and Erasmus MC for the GFP-expressing HMPV NL/1/00 strain.

