## Supplement for "Critical Nodes of Virus–Host Interaction Revealed Through an Integrated Network Analysis"

### Supplementary Material

**A)**

| Virus | Genome type | Number of baits | Experiment type | Number of interactions |
|---|---|---|---|---|
| Adeno-associated virus 5 (AAV5) | dsDNA | 7 | AP-MS and Y2H | 154 |
| Dengue virus (DENV) | ssRNA+ | 8 | Y2H | 135 |
| Epstein-Barr virus (EBV) | dsDNA | 56 | AP-MS and Y2H | 1887 |
| Influenza A virus PR8 (IAV-PR8) | ssRNA- | 11 | AP-MS and Y2H | 209 |
| Influenza virus Udorn (IAV-Udorn) | ssRNA- | 8 | AP-MS and Y2H | 95 |
| Hepatitis C virus (HCV) | ssRNA+ | 11 | Y2H | 466 |
| Human immunodeficiency virus 1(HIV-1) | ssRNA-RT | 16 | AP-MS | 514 |
| Human papilloma virus 5 (HPV5) | dsDNA | 2 | AP-MS and Y2H | 62 |
| Human papilloma virus 6B (HPV6B) | dsDNA | 4 | AP-MS and Y2H | 504 |
| Human papilloma virus 8 (HPV8) | dsDNA | 2 | AP-MS and Y2H | 158 |
| Human papilloma virus 11 (HPV11) | dsDNA | 4 | AP-MS and Y2H | 278 |
| Human papilloma virus 16(HPV16) | dsDNA | 5 | AP-MS and Y2H | 257 |
| Human papilloma virus 18 (HPV18) | dsDNA | 5 | Y2H | 368 |
| Human papilloma virus 33 (HPV33) | dsDNA | 3 | Y2H | 54 |
| Merkel cell polyomavirus (MCPyV) | dsDNA | 3 | AP-MS and Y2H | 104 |
| Simian virus 40 (SV40) | dsDNA | 2 | AP-MS and Y2H | 179 |
| Vaccinia virus (VACV) | dsDNA | 36 | AP-MS and Y2H | 357 |

**B)**

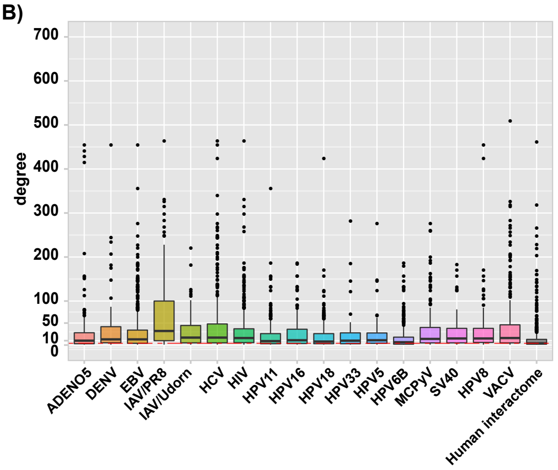


**C)**

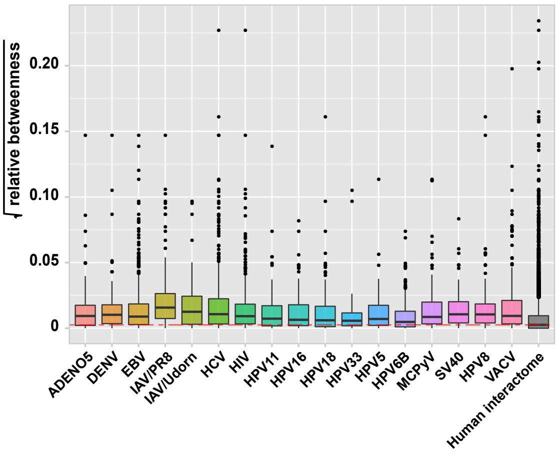


**Fig. S1: Properties of the hvPPI dataset.** (A) Table describing the details of the 17 viruses, the genome type, number of baits, experimental type and number of interactions with the host proteins. (B) and (C) Boxplots showing the distribution of degree and betweenness centrality of targets of each virus as compared to the human proteome.

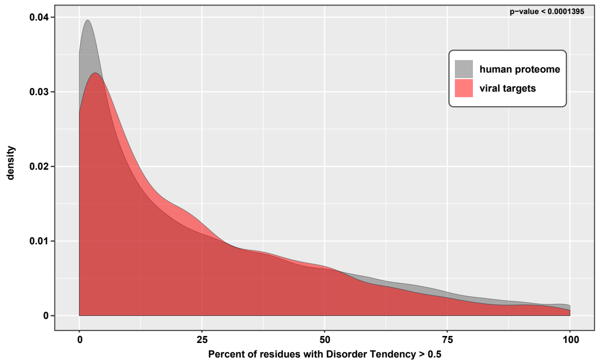


**Fig. S2: Protein disorder analysis.** Density plot of the distribution of host proteins in the hvPPI with percent of residues with disorder tendency greater than 0.5 as predicted by IUPred2A software as compared to the human proteome.

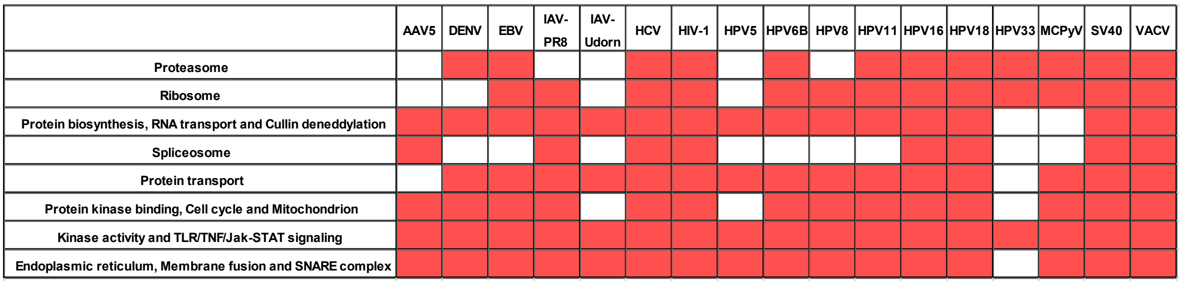


**Fig. S3: Clusters of hvPPI involved in core cellular processes.** Statistically significant cellular processes associated with the host proteins targeted by each virus. Red color shows statistically significant enrichment (p value $<$ 0.01) for that particular virus.

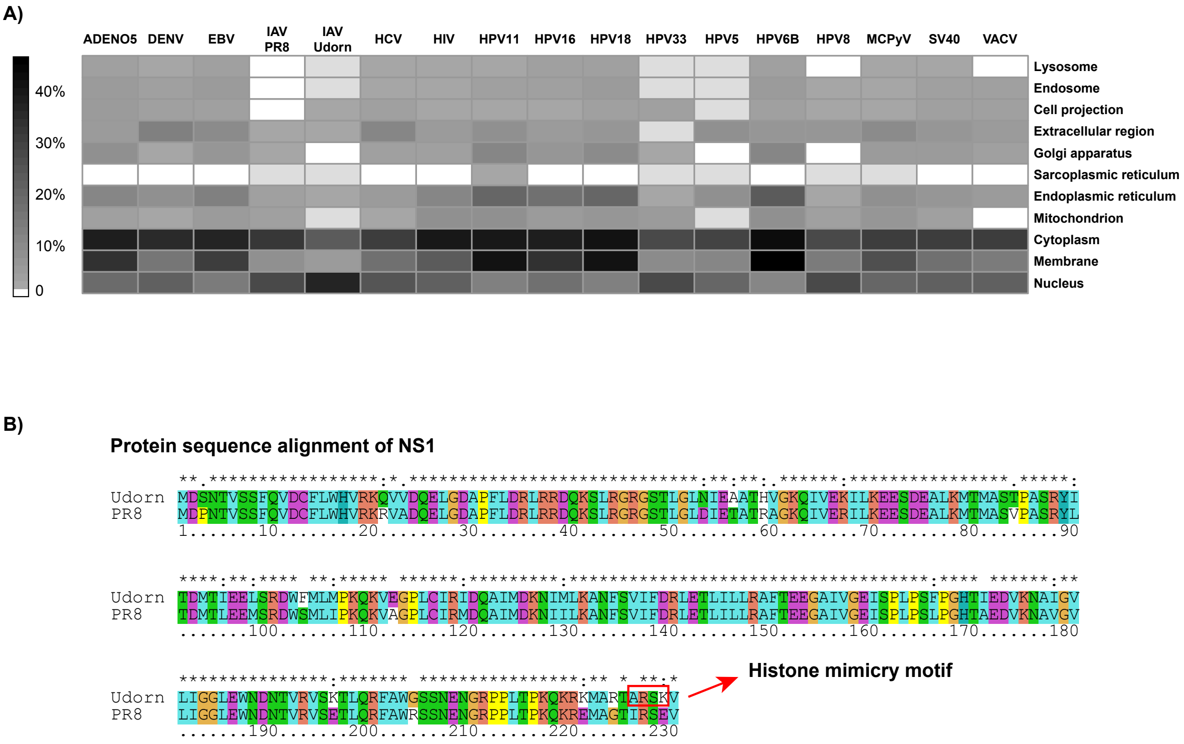


**Fig. S4: Sub-cellular localisation of the host proteins.** (A) Heatmap showing the sub-cellular location of the host proteins for the 17 viruses. (B) Protein sequence alignment of NS1 proteins from IAV strains PR8 and Udorn.

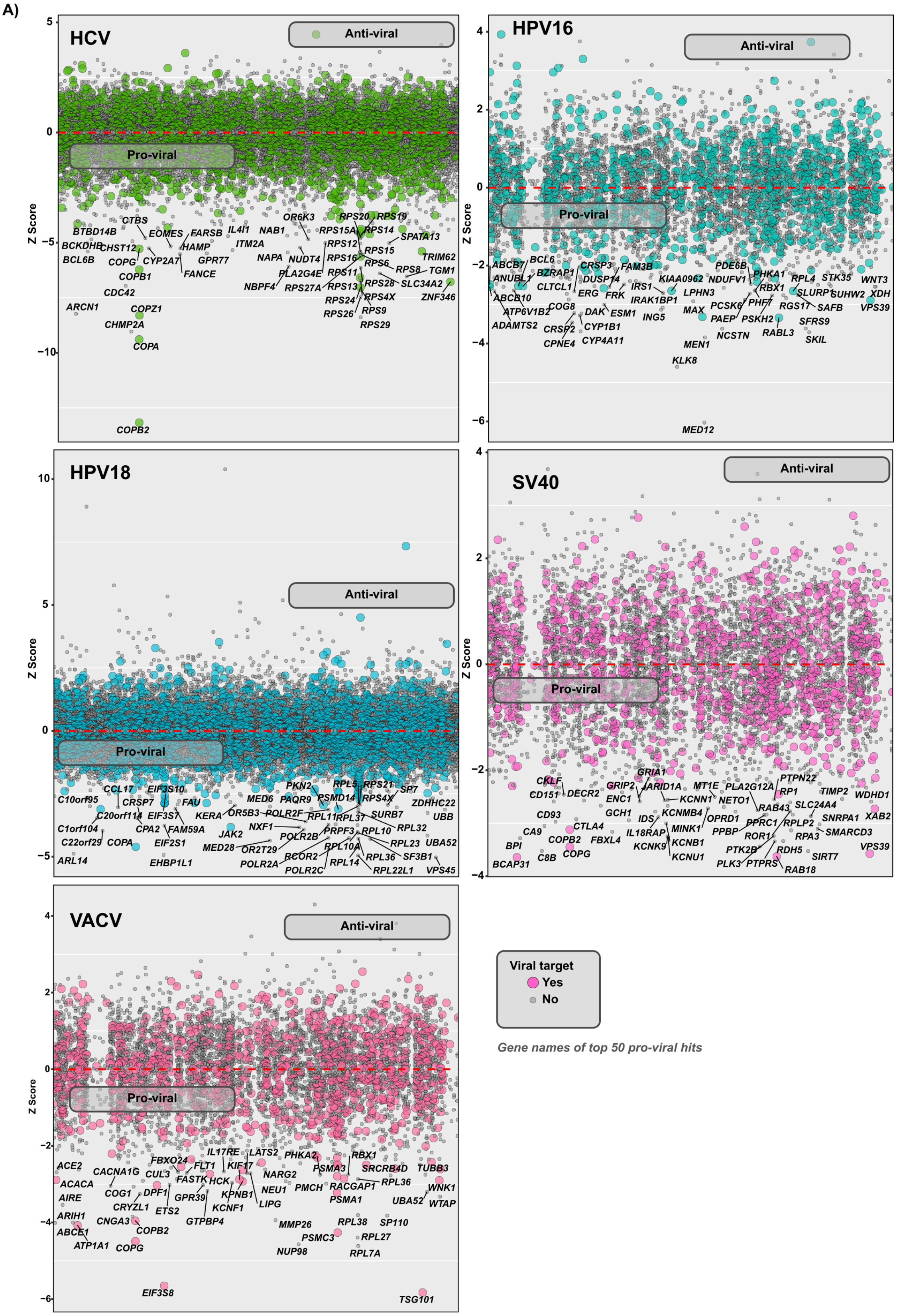


**Fig. S5: Integration of hvPPI with RNAi screen.** Scatter plot showing the viral replication phenotype for 5 different viruses. Gene names for the top 50 proviral genes are highlighted. Genes/Proteins which are part of the hvPPI are shown in bigger dots with colors.

**Table S1:** Drug re-purposing screen for novel broad-spectrum antivirals. Table showing the tested broad-spectrum antivirals and their representation in the hvPPI network.

| Broad–spectrum antiviral drug | # of targets in the host–virus network | # of host-virus interactions |
|---|---|---|
| ACYCLOVIR | 1 | 1 |
| ASPIRIN | 6 | 20 |
| AZACYTIDINE | 2 | 2 |
| AZITHROMYCIN | 1 | 7 |
| BORTEZOMIB | 47 | 219 |
| CAFFEINE | 2 | 2 |
| CYCLOSPORINE | 15 | 42 |
| DASATINIB | 14 | 37 |
| DIBUCAINE | 1 | 1 |
| ERLOTINIB | 4 | 16 |
| GEFITINIB | 3 | 11 |
| GEMCITABINE | 8 | 15 |
| HYDROXYCHLOROQUINE | 2 | 9 |
| IMATINIB | 7 | 26 |
| INDOMETHACIN | 3 | 4 |
| IVERMECTIN | 1 | 1 |
| LAMIVUDINE | 2 | 9 |
| LOVASTATIN | 5 | 21 |
| METFORMIN | 2 | 5 |
| MYCOPHENOLICACID | 3 | 28 |
| PENTOSANPOLYSULFATESODIUM | 1 | 1 |
| RIBAVIRIN | 1 | 7 |
| RITONAVIR | 1 | 7 |
| SIMVASTATIN | 1 | 1 |
| SIROLIMUS | 14 | 40 |
| TOPOTECAN | 4 | 16 |
| TRIFLURIDINE | 1 | 5 |
| VERAPAMIL | 3 | 12 |
| ITRACONAZOLE | 0 | 0 |
| ORITAVANCIN | 0 | 0 |
| LOPINAVIR | 0 | 0 |
| NITAZOXANIDE | 0 | 0 |
| CIDOFOVIR | 0 | 0 |
| MINOCYCLINE | 0 | 0 |
| PIRLINDOLEMESYLATE | 0 | 0 |
